# LUCID: joint cell typing and neighbour-aware decontamination of segmented spatial transcriptomics

**DOI:** 10.64898/2026.11.07.765042

**Authors:** Francesca Cupaioli, Ivan Merelli

## Abstract

Cell profiles in segmented spatial transcriptomics are contaminated by RNA of neighbouring cells and by segmentation errors that merge two cells. LUCID is a multinomial mixture model that jointly assigns a type to each cell, tests whether its profile mixes two types and estimates the fraction of counts coming from its neighbours. An external reference is adapted to the platform, an identifiability rule protects a cell’s own signal and the test is calibrated by parametric bootstrap. LUCID works with or without a reference, runs on CPUs and returns integer counts for all genes. On three public Visium HD datasets (mouse brain, human colorectal cancer, mouse liver) and in simulations built on them, LUCID removed contamination from other cell types as well as two-step pipelines, including RCTD followed by DecontX, flagged fewer contaminated cells as mixed, emptied no cell and preserved minority lineages where one cell type dominates.

**Availability and implementation:** LUCID is written in Python and is freely available under the MIT licence at https://www.bioinfotiget.it/gitlab/custom/lucid, with the scripts that reproduce all analyses from public data.

**Supplementary information:** Supplementary data are available online.

## Introduction

Spatial transcriptomics platforms with subcellular resolution, such as Visium HD, assign transcripts to segmented cells. Two artefacts make the resulting profiles impure. RNA from neighbouring cells and from the extracellular space is captured inside a cell’s boundary (ambient contamination), and segmentation errors merge two cells into one (doublets). Both produce profiles that mix more than one cell type and bias cell typing and differential expression.

Tools designed for droplet data, such as SoupX (Young and Behjati, 2020) and DecontX (Yang *et al*., 2020), estimate contamination from a global or cluster-level profile and ignore tissue position. RCTD (Cable *et al*., 2022) decomposes a profile into one or two cell types but has no ambient term, so contamination from the surroundings can be read as a second cell. ResolVI (Ergen and Yosef, 2026) models neighbour diffusion and background in a deep generative model trained on a GPU, typically on a few thousand highly variable genes (HVGs), and returns model-based expression estimates. Recent preprints address decontamination of spatial data with identifiability constraints (Wang and Gentles, 2026) and of single-cell genomics data in general (Caskey *et al*., 2026).

LUCID (Likelihood Unmixing of Cell Identity and Decon-tamination) addresses the two artefacts in one explicit like-lihood. For every cell it reports a cell type, a calibrated test for a mixed profile, the fraction of counts attributed to the surroundings, and a decontaminated integer count vector for all measured genes. It runs with a reference of cell-type pro-files or without one.

## methods

### Model

For cell *c* of sample *s* with *N*_*c*_ counts, LUCID compares two multinomial models of the count vector **y**_*c*_

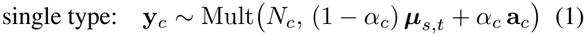

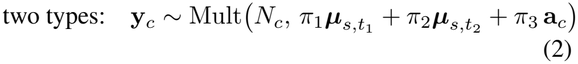

where ***µ***_*s,t*_ is the profile of type *t* in sample *s*, **a**_*c*_ is the ambient profile of the cell, *α*_*c*_ is its ambient fraction and ***π*** lies on the simplex. The ambient weight carries an exponential prior, a penalty *λα* on the log-likelihood (*λ* = 10), so that it shrinks to zero when the data do not support it. Type profiles are shared across samples up to a gene-wise sample factor.

### Ambient profile from the neighbours

Contamination is local. LUCID builds **a**_*c*_ from the cell’s *k* nearest segmented cells (*k* = 6): the count-weighted type composition of the neighbours is multiplied by the type profiles and mixed (weights 0.8 and 0.2) with a regional profile estimated from bins not assigned to any cell.

### Identifiability rule

When a cell is surrounded by cells of its own type, its ambient profile equals its own profile and *α*_*c*_ cannot be estimated. Without a safeguard, any mismatch between the reference and the tissue makes the ambient term absorb the whole cell. If the neighbours of a cell are in majority of one type, LUCID gives that type a second state for the cell, with **a**_*c*_ as profile and *α*_*c*_ = 0, and selects it when it fits at least as well as the reference state. LUCID thus removes RNA of other cell types and leaves RNA of the cell’s own type untouched.

### Platform adaptation of the reference

A single-cell reference differs from the spatial assay in the capture efficiency of each gene, as noted for RCTD (Cable *et al*., 2022). LU-CID fits the pooled profile of the segmented cells as a non-negative mixture of the reference types, with robust weights, and multiplies every type profile by the gene-wise ratio between the pooled profile and the fit. The factor is the same for all types, so it carries no information on cell identity. Genes absent from the reference start from the pooled pro-file.

### Fitting

All types are screened on a grid of *α*; the three best are refined by a safeguarded Newton method. Partner types for model (2) are chosen by a score test and the weights are fitted by EM. Type profiles are then re-estimated from confidently typed cells, with the adapted reference as prior. In reference-free mode the starting profiles are Leiden clusters (Traag *et al*., 2019) of the data.

### Calibrated test and output

The log-likelihood ratio between models (2) and (1) is compared with a null distribution obtained by simulating singlets from the fitted model, with estimated overdispersion. Cells with *p <* 0.05 are flagged as mixed profiles. Decontaminated counts are the posterior expected counts of the cell’s own (or major) type, rounded to integers by the largest-remainder rule so that totals are preserved. Details are given in Supplementary Methods.

### Implementation

LUCID is written in Python (NumPy, SciPy, pandas), reads and writes AnnData files (Virshup *et al*., 2024) and uses Scanpy (Wolf *et al*., 2018) for the reference-free initialisation. On three CPU cores, 39,854 cells and 19,070 genes took 37 min and 215,405 cells and 18,132 genes took 94 min.

## Results

### Simulation with spatial contamination

We placed 39,854 simulated cells of 49 brain types on the coordinates of the mouse brain section analysed below. Each cell received RNA from its ten nearest neighbours at a true ambient fraction of 0, 0.1, 0.3 or 0.5, and 6,004 cells were merged with a neighbour. The true profiles differed from the reference given to the methods by the gene-wise platform factor estimated on the real section and by random noise (Supplementary Note 1). As comparators representing the usual two-step design we used RCTD in doublet mode (Cable *et al*., 2022) followed by DecontX (Yang *et al*., 2020) on the RCTD types, and a sequential baseline with the same structure run with the adapted reference of LUCID (Supplementary Notes 1 and 6).

On the 2,000 HVGs used by ResolVI, the total-variation distance between the corrected and the true profile of the 33,850 singlets was 0.127 for raw counts, 0.134 for ResolVI (0.114 with RCTD labels), 0.052 for the sequential baseline, 0.071 for RCTD + DecontX, 0.061 for LUCID with reference and 0.058 for reference-free LUCID (Fig. 1B). The gain came from the 14,932 cells with an ambient fraction of at least 0.3 and neighbours mostly of other types (0.233 for raw counts, 0.214 and 0.178 for ResolVI without and with labels, 0.124 for RCTD + DecontX, 0.093 to 0.094 for the baseline and LUCID). For the 8,449 uncontaminated cells all methods stayed close to raw counts (0.068 to 0.075). On all genes the distance was 0.080 for raw counts, 0.064 for the baseline, 0.078 for RCTD + DecontX, 0.069 for reference-free LUCID and 0.081 for LUCID with reference.

**Figure 1.**
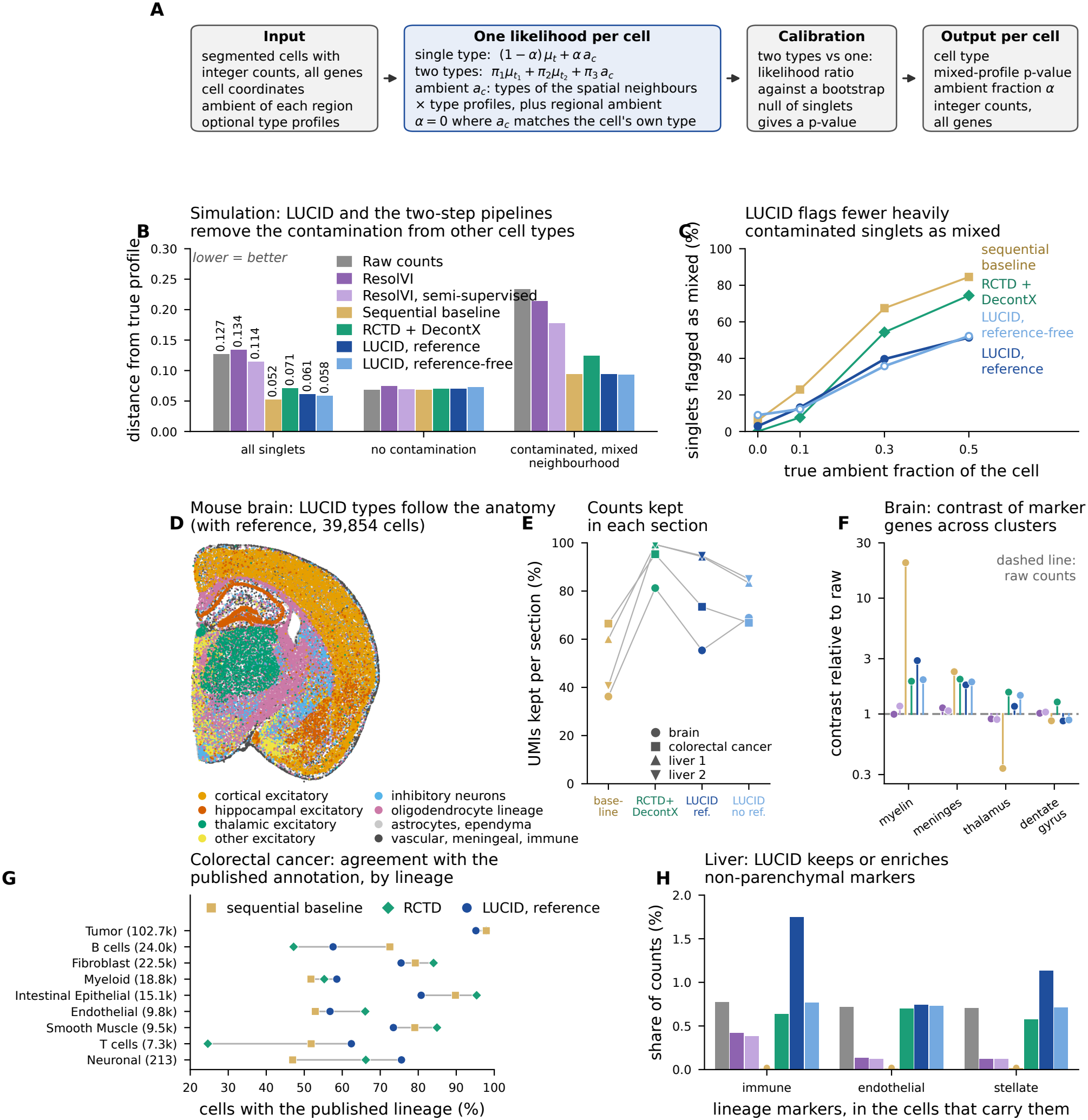
LUCID model and validation on public data. (**A**) Workflow. (**B**) Spatial simulation (39,854 cells). Total-variation distance between corrected and true profile on 2,000 HVGs, averaged over true singlets: all (*n* = 33,850), un-contaminated (*n* = 8,449), and contaminated with an ambient fraction of at least 0.3 and at least half of the neighbours of other types (*n* = 14,932). The sequential baseline is a two-type test without ambient term followed by regional ambient subtraction. RCTD + DecontX is RCTD in doublet mode followed by DecontX with default settings. In B, F and H, ResolVI is shown without labels and semi-supervised with RCTD labels. (**C**) Share of true singlets flagged as mixed by true ambient fraction. (**D**) LUCID types (with reference) in the mouse brain section, grouped in eight classes. (**E**) Share of counts kept in each of the four sections. (**F**) Contrast of marker genes between the Space Ranger cluster where they are expected and the clusters where they are not, on the HVGs, relative to raw counts (log scale). (**G**) Colorectal cancer: share of cells whose lineage agrees with the published unsupervised annotation; cells per lineage in brackets. RCTD: first RCTD type. (**H**) Mean share of lineage-marker counts, on the HVGs, in liver cells with at least three immune (*n* = 4,100), endothelial (*n* = 543) or stellate (*n* = 557) marker counts in the raw data. Colours as in B.

The two-step pipelines reached this result by treating contamination as a second cell: the baseline and RCTD called 45.2% and 34.0% of the true singlets doublets, and 84.5% and 74.4% of those with an ambient fraction of 0.5. LU-CID flagged 26.7% with reference and 27.3% without, and 2.9% and 9.0% of the uncontaminated singlets (Fig. 1C). It assigned the correct type to 97.0% of the singlets with reference (baseline 95.3%, RCTD 91.8%). A cell merged with a neighbour and a cell heavily contaminated by that neighbour have the same profile, and LUCID detected 55% and 62% of the merged cells (baseline 87%, RCTD 79.8%). In a second simulation with region-level contamination, where the two cases differ, LUCID with reference flagged 6.8% of singlets and detected 81% of the merged cells, against 47.6% and 95% for the baseline and 35.3% and 85.2% for RCTD (Supplementary Notes 1 and 6).

### Mouse brain

A coronal section of mouse brain from 10x Genomics, segmented by Space Ranger (39,854 cells), was analysed with 49 types from the Allen whole-brain atlas (Yao *et al*., 2023). An atlas from another technology is the hardest case for a reference: the platform factor had a standard deviation of 0.92 on the natural-log scale, and without adaptation LUCID kept 31.4% of the counts (Supplementary Note 5). With adaptation, types followed the anatomy (Fig. 1D): 91% of layer 2/3 intratelencephalic neurons lay in the cortical niche (34% of all cells), 82% of thalamic glutamatergic neurons in the thalamic niche (8%) and 88% of dentate granule cells in the dentate niche (3%).

LUCID kept 55.4% of the counts with reference and 68.9% without, and emptied no cell and one cell. The baseline kept 36.2% and emptied 3,790 cells, RCTD + DecontX kept 81.2% and emptied 458 (Fig. 1E). As a model-free check we computed, for marker genes of four structures, the ratio between their share of counts in the Space Ranger cluster where they are expected and in the clusters where raw counts show less than a tenth of that share. On the HVGs, LUCID raised the contrast for myelin (2.89- and 1.98-fold, with and without reference), meninges (1.78 and 1.89) and thalamus (1.16 and 1.45) and lowered it for the dentate gyrus (0.87 and 0.89). ResolVI stayed between 0.91 and 1.13 times the raw contrast (0.90 to 1.17 with labels), the baseline ranged from 0.34 for thalamus to 20 for myelin and RCTD + DecontX raised all four (1.27 to 2.00, Fig. 1F). On all genes the dentate contrast rose with LUCID and the meningeal contrast fell with reference (Supplementary Note 2). LUCID flagged 43.6% (with reference) and 31.5% (reference-free) of the cells as mixed, RCTD called 76.9% doublets.

### Human colorectal cancer

A colorectal cancer section of Oliveira *et al*. (2025) (215,405 cells) was analysed with the 33 cell types of the single-cell reference of the same study. Here the reference already matched the platform (standard deviation of the log factor 0.39) and adaptation changed little. LUCID kept 73.5% of the counts with reference and 66.8% without and emptied at most one cell. The base-line kept 66.5% and emptied 5,890 cells, RCTD + DecontX 95.3% and 23. The lineage of the LUCID type agreed with the published unsupervised annotation for 80.5% of the cells, against 83.7% for the baseline and 82.0% for RCTD. The two published annotations agree with each other for 84.7%. The two-step pipelines agreed more often for the abundant lineages, LUCID for T cells (62.4%, against 51.8% and 24.6%), myeloid and neuronal cells (Fig. 1G). Agreement fell from 94% for cells with an estimated ambient fraction below 0.05 to 56% above 0.6, so *α*_*c*_ marks the cells whose identity is uncertain. On the HVGs, LUCID raised the contrast of T-cell, fibroblast and B-cell markers between their published lineage and the others (for T cells 19.6 against 10.9 for raw counts), ResolVI did not (10.3, and 11.2 with labels) and RCTD + DecontX raised it more (Supplementary Note 6). For epithelial markers LUCID with reference lowered it (37.8 against 46.5, Supplementary Note 3). LUCID flagged 39.4% of the cells as mixed, the baseline 60.9% and RCTD 31.8%.

### Mouse liver

The liver is a stress test, because one cell type dominates and the ambient profile coincides with the hepatocyte profile. In two replicate sections segmented by the authors (Novoselsky and Itzkovitz, 2026) (49,331 cells), with 8 types from Tabula Muris Senis (Tabula Muris Consortium, 2020), LUCID kept 94.5% of the counts with reference and 84.4% without, and assigned 91.8% of the cells to hepatocytes, 4.1% to sinusoidal endothelium and 3.2% to stellate cells. The baseline kept 60.0% and 40.7% of the counts in the two replicates, emptied 468 cells and typed 99.9% of the cells as hepatocytes. RCTD typed 99.9% as hepatocytes and DecontX kept 99.3% of the counts. In cells carrying immune, endothelial or stellate marker counts, the share of those markers on the HVGs rose or held with LUCID with reference (0.77% to 1.74%, 0.71% to 0.74% and 0.71% to 1.13%), did not change without reference, fell with ResolVI with or without labels, fell or held with RCTD + DecontX and vanished with the baseline (Fig. 1H). The pericentral to periportal gradient of zonation markers along the published lobule zones was unchanged by LUCID (Supplementary Note 4).

## Limitations

LUCID cannot tell a cell merged with a neighbour from a cell contaminated by that neighbour, so its flag denotes a mixed profile and not a doublet. Given the same adapted reference, the sequential baseline matched or exceeded LUCID on profile distance in simulation, both two-step pipelines agreed more often with the published annotation of the tumour, and RCTD + DecontX gave a higher marker contrast in most brain structures and tumour lineages. This contrast rewards removing counts outside a cell’s cluster, not fidelity to the true profile, which in simulation was higher with LU-CID. The advantages of LUCID are a more specific flag, no emptied cells and the behaviour where one cell type dominates. On real data the bootstrap null was permissive: at the nominal 5% it flagged 6.7% of hepatocytes without lineage-marker counts, 8.1% of cells in a white-matter cluster of the brain (reference-free) and 22.7% of tumour cells in the tumour core. With the atlas reference LUCID removed almost half of the brain counts and placed 71% of its 1,043 hippocampal CA3 cells in the cortical niche. RNA of a cell’s own type coming from neighbours is not removed. The reference-free mode depends on the initial clustering. Each tissue is represented by one section or two replicates, so we make no claim on differential expression.

## Acknowledgements

We thank the authors of the public datasets and the CNR-ITB HPC infrastructure.

## Conflict of Interest

None declared.

## Data Availability

All data are public. Mouse brain and human colorectal cancer: 10x Genomics datasets Visium_HD_Mouse_Brain and Visium_HD_Human_Colon_Cancer (Space Ranger 4.0.1 outputs with cell segmentation, CC BY 4.0), downloaded from https://cf.10xgenomics.com/samples/spatial-exp/4.0.1/. Colorectal cancer raw data: GEO GSE280318. Its annotations and single-cell reference: https://github.com/10XGenomics/HumanColonCancer_VisiumHD. Mouse liver: Zenodo, https://doi.org/10.5281/zenodo.17779250. Brain reference: Allen Brain Cell Atlas, WMB-10Xv3 expression matrices (release 20230630). Liver reference: Tabula Muris Senis on CEL-LxGENE. Source code and analysis scripts are available at https://www.bioinfotiget.it/gitlab/custom/lucid.

## Notes

### Competing Interest Statement

The authors have declared no competing interest.

### Summary of Updates

Added a benchmark against two-step pipelines (RCTD + DecontX, default and background modes) and a semi-supervised ResolVI run, across all datasets and simulations. Figure 1 and Figures S1-S2 updated with RCTD+DecontX; new Supplementary Note 6 with Tables S17-S24 and the Cable (2022) and Yang (2020) references. Title unchanged; abstract edited in its final sentence only; still a 4-page Application Note.

