## Supplementary_File for "LUCID: joint cell typing and neighbour-aware decontamination of segmented spatial transcriptomics"

### Supplementary Material for:
### LUCID: joint cell typing and neighbour-aware decontamination of segmented spatial transcriptomics

Francesca Cupaioli and Ivan Merelli

### Supplementary Methods

#### *Notation and likelihood*

For cell $c$ of sample $s$, $\mathbf{y}_c \in \mathbb{Z}_{\geq 0}^G$ is the count vector over the $G$ measured genes and $N_c = \sum_g y_{cg}$. Type profiles $\boldsymbol{\mu}_t$ ($t = 1, \ldots, T$) and ambient profiles $\mathbf{a}_c$ are points of the simplex. The profile of type $t$ in sample $s$ is $\boldsymbol{\mu}_{s,t} \propto \boldsymbol{\mu}_t \, \boldsymbol{\gamma}_s$, where $\boldsymbol{\gamma}_s$ is a gene-wise sample factor. The two models are

$$\ell_1(t, \alpha) = \sum_g y_{cg} \log\big[(1-\alpha)\mu_{s,t,g} + \alpha a_{cg}\big] - \lambda\alpha, \tag{S1}$$

$$\ell_2(t_1, t_2, \boldsymbol{\pi}) = \sum_g y_{cg} \log\big[\pi_1 \mu_{s,t_1,g} + \pi_2 \mu_{s,t_2,g} + \pi_3 a_{cg}\big] - \lambda\pi_3, \tag{S2}$$

with $\alpha \in [0, 1)$ and $\boldsymbol{\pi}$ on the simplex. The term $\lambda\alpha$ is an exponential prior on the ambient weight; $\lambda = 10$ in all analyses.

#### *Platform adaptation of the reference*

A single-cell reference and a spatial assay capture the same transcript with different efficiency (chemistry, probe design, fixation). LUCID removes this difference before fitting, in two steps applied to the reference restricted to the genes of the spatial data. First, genes with no count in the reference start from the pooled profile of the segmented cells, the same value for every type, and the other genes are rescaled so that each profile sums to one; these genes then carry no information on the type until the profiles are re-estimated. Second, the pooled profile $\mathbf{s}$ of the segmented cells is fitted as a non-negative mixture of the reference profiles, $\mathbf{s} \approx \sum_t w_t \boldsymbol{\mu}_t$, by non-negative least squares on the genes weighted by $1/\sqrt{s_g}$ and by Huber weights on the log residual (threshold 1, 15 iterations). The platform factor of gene $g$ is $f_g = s_g / \sum_t w_t \mu_{t,g}$, clipped to $[1/20, 20]$, and every profile is multiplied by $\mathbf{f}$ and renormalised. The factor is shared by all types: it changes how well a gene is captured, not which type expresses it. In a test on simulated data the factor was recovered with a correlation of 0.997 on the log scale. The step is the default of the command-line driver and can be switched off. The same adapted reference was given to the sequential baseline.

#### *Fitting*

*Screening.* For every cell and type, $\ell_1$ is evaluated on the grid $\alpha \in \{0.02, 0.15, 0.35, 0.6, 0.85\}$. The three types with the highest value are refined: $\alpha$ is found by Newton iterations on the concave penalised likelihood, safeguarded by bisection (25 iterations), and values below $10^{-4}$ are set to zero. The type with the highest $\ell_1$ is the singlet type $\hat{t}$ with ambient fraction $\hat{\alpha}$.

*Partner types.* Given the fitted singlet mixture $\mathbf{m}_c$, the score $\sum_g y_{cg} \, \mu_{s,t,g}/m_{cg} - N_c$ measures how much the likelihood would gain by adding type $t$. The two types with the highest score $(u_1, u_2)$ define three candidate pairs, $(\hat{t}, u_1)$, $(\hat{t}, u_2)$ and $(u_1, u_2)$. For each pair the weights $\boldsymbol{\pi}$ are fitted by EM with the closed-form penalised M-step (40 iterations). The test statistic is $\Lambda_c = \max\big(0, \max_{\text{pairs}} \ell_2 - \ell_1(\hat{t}, \hat{\alpha})\big)$.

*Reference refinement.* In two rounds, cells whose margin between the best and the second-best type (per count) is above the median of their type, and whose $\hat{\alpha} \leq 0.7$, are used to re-estimate the type profiles from the posterior counts of the cell's own type. The adapted reference enters as a prior worth $2 \times 10^4$ counts per type. With more than one sample, the gene-wise factor $\boldsymbol{\gamma}_s$ is estimated from cells of the dominant type in groups whose regional ambient is close to that type (cosine at least 0.9), shrunk towards one with a prior worth 20% of the counts and clipped to $[0.2, 5]$.

*Reference-free mode.* Counts are normalised, log-transformed and reduced to 30 principal components on

2,000 highly variable genes; Leiden clustering (Traag *et al.*, 2019) at resolution 1.0 on a 15-neighbour graph gives the starting profiles (clusters of at least 50 cells), which are then refined as above.

#### *Ambient profile*

*Regional ambient.* For every sample and tissue region a profile $\mathbf{r}$ is obtained from the Visium HD bins that are not assigned to any segmented cell. During reference refinement $\mathbf{a}_c = \mathbf{r}_{r(c)}$.

*Local ambient.* After refinement, each cell is linked to its $k$ nearest segmented cells of the same section ($k = 6$), discarding neighbours farther than three times the median nearest-neighbour distance. With $n_c(t)$ the count-weighted share of type $t$ among the neighbours,

$$\mathbf{a}_c = \beta \sum_t n_c(t)\, \boldsymbol{\mu}_{s,t} + (1-\beta)\, \mathbf{r}_{r(c)}, \qquad \beta = 0.8. \tag{S3}$$

Cells without neighbours keep the regional profile. We build the local profile from the type composition of the neighbours and not from their pooled counts, because pooled counts of a few cells are a noisy estimate of the contamination profile and inflate the test statistic.

#### *Identifiability rule*

If a cell of type $t$ is surrounded by cells of type $t$, then $\mathbf{a}_c \approx \boldsymbol{\mu}_{s,t}$ and $\alpha_c$ is not identifiable. In practice the measured ambient matches the tissue better than an external reference does, and the unconstrained model assigns the whole cell to the ambient. LUCID applies the following rule. During refinement, a region is flagged when the cosine between its ambient and the pooled profile of its cells is at least 0.99 and one type explains at least half of the ambient in a mixture decomposition. In the local stage, a cell is flagged when one type accounts for at least half of its neighbourhood. For a flagged cell that type has a second state with profile $\mathbf{a}_c$ and $\alpha = 0$, which is selected when its likelihood is at least that of the reference state. The consequence is that LUCID does not remove RNA of the cell's own type.

#### *Calibration of the mixed-profile test*

A random subset of 20,000 cells is re-simulated as singlets from the fitted model, each with its own $\hat{t}$, $\hat{\alpha}$, $\mathbf{a}_c$ and $N_c$. Counts follow a Dirichlet-multinomial distribution whose concentration is estimated by the method of moments on the 200 most expressed genes of the most frequent type. The simulated cells are analysed with the same procedure and their $\Lambda$ values give the null distribution, separately for each sample-by-region group with at least 300 simulated cells. Cells with $p < 0.05$ are reported as mixed and those with $p < 0.01$ as high-confidence mixed.

#### *Decontaminated counts*

For a single-type cell the share of gene $g$ that belongs to the cell is $\rho_{cg} = (1-\hat{\alpha})\mu_{s,\hat{t},g}/[(1-\hat{\alpha})\mu_{s,\hat{t},g} + \hat{\alpha} a_{cg}]$; for a mixed cell it is the posterior share of the type with the larger weight. The expected counts $y_{cg}\rho_{cg}$ are rounded to integers by the largest-remainder rule, so that the total of each cell equals the rounded expected total.

#### *Sequential baseline*

The baseline reproduces the common two-step design. Cells are typed by maximum multinomial likelihood on the reference profiles without ambient term. A two-type mixture is fitted for the three most likely types and a cell is called a doublet when the log-likelihood ratio exceeds fixed thresholds (25 for certain, 4 for uncertain doublets), after which the second-type share is subtracted. A per-cell ambient fraction is then estimated against the regional ambient profile by EM (30 iterations, capped at 0.95) and removed. The baseline received the platform-adapted reference, as LUCID did. With the reference as given it kept 14.1% of the counts in the brain, 53.2% in the colorectal cancer and 11.3% and 6.8% in the two liver replicates.

#### *ResolVI*

ResolVI (Ergen and Yosef, 2026) was run with scvi-tools 1.4.0 (Gayoso *et al.*, 2022) in unsupervised mode on the 2,000 highly variable genes selected by Scanpy with the section as batch, with spatial coordinates, 100 epochs and a batch size of 1,024 on one GPU. The corrected expression is the mean of five posterior samples of the normalised expression. ResolVI was run on the three sections and on the spatial simulation; the jobs took 5 (brain), 7 (liver), 22 (colorectal cancer) and 8 min (simulation). ResolVI requires at least five counts per cell, so 6,669 cells of the colorectal cancer section with fewer counts on the HVGs were left out and counted as empty. ResolVI was also run in its semi-supervised mode on the same cells and genes, with the first RCTD type of each cell as label and the cells rejected by RCTD left unlabelled; training took 7, 9, 36 and 7 min (Supplementary Note 6).

***Public datasets and preparation***

*Mouse brain.* Visium HD Spatial Gene Expression of a formalin-fixed coronal section of mouse brain, distributed by 10x Genomics with Space Ranger 4.0.1 outputs that include cell segmentation. *Human colorectal cancer.* Visium HD section of patient P2 of Oliveira *et al.* (2025), distributed by 10x Genomics with Space Ranger 4.0.1 segmentation. The published annotations of the 8 $\mu$m bins (unsupervised clusters, deconvolution labels and classes, tumour periphery regions) were transferred to the segmented cells by position. *Mouse liver.* Two replicate Visium HD sections of adult mouse liver (Novoselsky and Itzkovitz, 2026), with the cell masks, the lobule zone (1 pericentral to 6 periportal) and the segmentation class provided by the authors.

Cells with fewer than 100 counts were discarded (Table S1). The regional ambient profile is the sum of the counts of the tissue-covered 2 µm bins that are not assigned to any cell. In the brain and in the tumour the tissue was divided into eight niches by k-means on the composition of the 30 nearest cells in Space Ranger graph clusters, each bin was attributed to the niche of the nearest cell and one ambient profile was computed per niche. In the liver one profile was computed per replicate.

*References.* Brain: raw counts of the 10x v3 single-cell data of the Allen whole-brain atlas (Yao *et al.*, 2023) for the dissection regions covered by the section, aggregated to 49 types (atlas classes, with the cortical, hippocampal, glial, vascular and immune classes split into their main subclasses; at least 1,000 cells per type). Colorectal cancer: the Chromium Flex single-cell data of Oliveira *et al.* (2025), 33 types of the published level-2 annotation; lineages are the level-1 annotation of the same data. Liver: Tabula Muris Senis droplet data (Tabula Muris Consortium, 2020), 8 types with at least 30 cells.

Table S1: **Public sections.** Cells kept have at least 100 counts. UMIs in cells: share of the counts under tissue that fall inside a segmented cell.

| Section | Segmented cells | Cells kept | Genes | Median UMIs | UMIs in cells (%) | Ambient groups | Reference types |
|---|---|---|---|---|---|---|---|
| Mouse brain | 40,222 | 39,854 | 19,070 | 1,087 | 68.4 | 8 niches | 49 |
| Colorectal cancer | 220,703 | 215,405 | 18,132 | 780 | 88.7 | 8 niches | 33 |
| Mouse liver, replicate 1 | 24,578 | 20,873 | 14,703 | 1,028 | 52.2 | 1 | 8 |
| Mouse liver, replicate 2 | 33,785 | 28,458 | 14,798 | 1,710 | 74.6 | 1 | 8 |

***Software and run times***

LUCID was run with Python 3.12, SciPy 1.16.2, pandas 2.3.3, Scanpy 1.11.4 and anndata 0.13.4 on three CPU cores. Wall-clock times, including file reading and the bootstrap, are in Table S6. The released code (`lucid.py`, `lucid_run.py`) reproduced the decontaminated matrices of the liver analysis exactly (0 of 34,530,953 non-zero entries differed), both from the full reference and from a reference restricted to its own genes.

### Supplementary Note 1 — Simulations on public data

Both simulations use the mouse brain section and its reference. The reference given to the methods is the atlas reference. The true profiles are the atlas profiles multiplied by the platform factor estimated on the real section and by log-normal noise specific to gene and type (standard deviation 0.3 in the regional and 0.1 in the spatial simulation). The type composition of each niche follows the LUCID typing of the real section (types above 2% of a niche). Counts are Dirichlet-multinomial with the overdispersion estimated on the real data. Each method adapts the reference on the simulated counts, as on real data.

***Simulation with regional contamination***

36,000 cells in eight niches, 15.2% true doublets (weight of the major cell between 0.3 and 0.7), true ambient fraction of 0, 0.1, 0.3 or 0.5 from the regional ambient profile, and 500, 1,000 or 2,000 counts per cell. Here a contaminated singlet and a merged cell have different profiles. LUCID with reference flagged 6.8% of the singlets and detected 80.9% of the doublets, with a precision of 0.68; the baseline flagged 47.6% of the singlets (Table S2, Fig. S1). Reference-free LUCID started from 21 clusters for 49 types and gave the type of the best-matching cluster to 73.7% of the singlets.

Table S2: **Simulation with regional contamination** (36,000 cells). Type acc.: singlets with the correct type (%). Flagged: cells flagged as mixed (%). Singlets fl.: true singlets flagged (%). Merged det.: true doublets flagged (%). The last four columns give the share of singlets flagged (%) by true ambient fraction.

| Method | Type acc. | Flagged | Singlets fl. | Merged det. | Precision | $\alpha$ MAE | 0 | 0.1 | 0.3 | 0.5 |
|---|---|---|---|---|---|---|---|---|---|---|
| Sequential baseline | 95.4 | 54.7 | 47.6 | 94.7 | 0.26 | 0.044 | 5.1 | 22.7 | 72.3 | 89.4 |
| LUCID, reference, $p < 0.05$ | 99.9 | 18.1 | 6.8 | 80.9 | 0.68 | 0.031 | 1.4 | 4.6 | 10.2 | 10.9 |
| LUCID, reference, $p < 0.01$ | 99.9 | 12.7 | 1.8 | 73.4 | 0.88 | 0.031 | 0.2 | 1.0 | 2.9 | 3.0 |
| LUCID, reference-free, $p < 0.05$ | 73.7 | 20.9 | 11.4 | 73.8 | 0.54 | 0.050 | 8.4 | 10.1 | 13.6 | 13.2 |
| LUCID, reference-free, $p < 0.01$ | 73.7 | 15.8 | 6.5 | 67.7 | 0.65 | 0.050 | 6.7 | 6.6 | 7.1 | 5.7 |

***Simulation with contamination from the neighbours***

39,854 cells on the coordinates of the real section; 6,004 cells were merged with a neighbour (weight between 0.15 and 0.85). Every cell received the mean true profile of its ten nearest neighbours at a true ambient fraction of 0, 0.1, 0.3 or 0.5, with 300 to 2,000 counts. Profile accuracy is the total-variation distance between the mean normalised corrected profile of the singlets of a type and the true profile of the type, averaged over types with the number of cells as weight (Table S3); the unweighted average over types is also given. With the same adapted reference the sequential baseline was as close to the truth as LUCID, or closer, because its two-type step subtracts the contaminating type as if it were a second cell. The price is the flag: the baseline called 51.4% of all cells doublets, LUCID 31.0% (Table S4). On all genes LUCID with reference did not improve on raw counts. Clustering of the ResolVI latent space gave an adjusted Rand index of 0.58 with the true types (0.49 for raw counts, 0.91 for the LUCID types with reference, 0.70 reference-free, 0.91 for the baseline).

Table S3: **Spatial simulation: distance from the true profile** of the true singlets ($n = 33{,}850$; uncontaminated $n = 8{,}449$; contaminated with ambient fraction of at least 0.3 and at least half of the neighbours of other types $n = 14{,}932$). Lower is better.

| Genes | Method | All singlets | Uncontaminated | Contaminated, mixed | All, macro average |
|---|---|---|---|---|---|
| 2,000 HVGs | Raw counts | 0.127 | 0.068 | 0.233 | 0.153 |
| 2,000 HVGs | ResolVI | 0.134 | 0.075 | 0.214 | 0.177 |
| 2,000 HVGs | Sequential baseline | 0.052 | 0.068 | 0.093 | 0.082 |
| 2,000 HVGs | LUCID, reference | 0.061 | 0.070 | 0.094 | 0.092 |
| 2,000 HVGs | LUCID, reference-free | 0.058 | 0.072 | 0.093 | 0.105 |
| All genes | Raw counts | 0.080 | 0.086 | 0.140 | 0.105 |
| All genes | Sequential baseline | 0.064 | 0.086 | 0.116 | 0.097 |
| All genes | LUCID, reference | 0.081 | 0.087 | 0.133 | 0.116 |
| All genes | LUCID, reference-free | 0.069 | 0.088 | 0.114 | 0.112 |

### Supplementary Note 2 — Mouse brain

Results per section and method are in Table S6. The 20 most frequent LUCID types by niche are in Table S7. The niches were named from their dominant types and position: cortex (N1), a mixed niche with striatum and hippocampal fields (N2), white matter (N3), hypothalamus (N4), thalamus (N5), meninges (N6) and dentate gyrus (N8). Two errors of the typing with the atlas reference are visible: 71% of the 1,043 cells typed as CA3 neurons lie in the cortical niche, in its ventral part, and only 53% of the 5,659

Table S4: **Spatial simulation: typing and flags.** Columns as in Table S2.

| Method | Type acc. | Flagged | Singlets fl. | Merged det. | Precision | $\alpha$ MAE | 0 | 0.1 | 0.3 | 0.5 |
|---|---|---|---|---|---|---|---|---|---|---|
| Sequential baseline | 95.3 | 51.4 | 45.2 | 86.6 | 0.25 | 0.071 | 6.1 | 22.9 | 67.5 | 84.5 |
| LUCID, reference, $p < 0.05$ | 97.0 | 31.0 | 26.7 | 55.3 | 0.27 | 0.079 | 2.9 | 13.1 | 39.6 | 51.4 |
| LUCID, reference, $p < 0.01$ | 97.0 | 19.3 | 15.2 | 42.9 | 0.33 | 0.075 | 1.0 | 4.2 | 21.8 | 33.7 |
| LUCID, reference-free, $p < 0.05$ | 72.5 | 32.5 | 27.3 | 62.0 | 0.29 | 0.106 | 9.0 | 12.2 | 35.8 | 52.2 |

Table S5: **Spatial simulation: number of neighbours used for the ambient profile.** The cells were contaminated by ten neighbours. Regional only: no local ambient.

| Method | Neighbours | Distance, HVGs | Distance, all genes | Singlets flagged | Merged detected | Type accuracy |
|---|---|---|---|---|---|---|
| LUCID, reference | regional only | 0.072 | 0.089 | 26.7 | 64.0 | 97.0 |
| LUCID, reference | 5 | 0.059 | 0.078 | 30.2 | 54.8 | 96.5 |
| LUCID, reference | 6 | 0.061 | 0.081 | 26.7 | 55.3 | 97.0 |
| LUCID, reference | 10 | 0.067 | 0.087 | 15.7 | 54.6 | 97.8 |
| LUCID, reference-free | regional only | 0.063 | 0.073 | 31.8 | 71.0 | 71.9 |
| LUCID, reference-free | 5 | 0.058 | 0.068 | 29.6 | 60.6 | 72.2 |
| LUCID, reference-free | 6 | 0.058 | 0.069 | 27.3 | 62.0 | 72.5 |
| LUCID, reference-free | 10 | 0.061 | 0.072 | 19.6 | 63.5 | 72.7 |

oligodendrocytes lie in the white-matter niche (13% of the cells), as expected from their presence in grey matter.

Table S6: **Results per section.** Emptied cells: cells left with no count. Minutes: wall-clock time on three CPU cores (one run for the two liver replicates).

| Section | Method | UMIs kept (%) | Mixed (%) | Emptied cells | Types or clusters | Minutes |
|---|---|---|---|---|---|---|
| Mouse brain | Sequential baseline | 36.2 | 90.2 | 3,790 | – | – |
| Mouse brain | LUCID, reference | 55.4 | 43.6 | 0 | 49 | 37 |
| Mouse brain | LUCID, reference-free | 68.9 | 31.5 | 1 | 14 | 26 |
| Colorectal cancer | Sequential baseline | 66.5 | 60.9 | 5,890 | – | – |
| Colorectal cancer | LUCID, reference | 73.5 | 39.4 | 0 | 33 | 94 |
| Colorectal cancer | LUCID, reference-free | 66.8 | 35.0 | 1 | 30 | 94 |
| Mouse liver, replicate 1 | Sequential baseline | 60.0 | 62.3 | 160 | – | – |
| Mouse liver, replicate 1 | LUCID, reference | 94.3 | 14.6 | 0 | 8 | 15 |
| Mouse liver, replicate 1 | LUCID, reference-free | 83.4 | 21.2 | 0 | 7 | 19 |
| Mouse liver, replicate 2 | Sequential baseline | 40.7 | 67.8 | 308 | – | – |
| Mouse liver, replicate 2 | LUCID, reference | 94.6 | 15.4 | 0 | 8 | 15 |
| Mouse liver, replicate 2 | LUCID, reference-free | 85.0 | 16.1 | 0 | 7 | 19 |

***Contrast of marker genes***

Marker sets: myelin (*Mbp, Plp1, Mobp, Mal, Mog, Mag, Cnp, Cldn11*), meninges (*Col1a1, Col1a2, Col3a1, Dcn, Lum, Igf2*), thalamus (*Tcf7l2, Prkcd, Slc17a6, Synpo2, Rgs16, Ntng1*), dentate gyrus (*Prox1, C1ql2, Dsp, Calb1, Pdzd2*) and cortical excitatory neurons (*Slc17a7, Satb2, Neurod6, Tbr1, Cux2, Rorb, Fezf2*; none among the HVGs). Groups are the Space Ranger graph clusters, which do not depend on any method compared here. For each set, the on-target cluster is the cluster of at least 300 cells with the highest mean share of marker counts in the raw data and the off-target clusters are those where the raw share is below a tenth of it. The contrast is the ratio between the mean share on target and off target (Table S8). The metric rewards any removal of counts off target, so it rises with aggressive subtraction: the baseline, which emptied 3,790 cells, has the highest myelin contrast and the lowest thalamic contrast. For LUCID the direction for meninges and dentate gyrus depends on the gene set.

Table S7: **Mouse brain: cells per LUCID type (with reference) and niche**, 20 most frequent types.

| Type | N1 | N2 | N3 | N4 | N5 | N6 | N7 | N8 | Total |
|---|---|---|---|---|---|---|---|---|---|
| Oligo NN | 676 | 673 | 2,971 | 450 | 532 | 20 | 321 | 16 | 5,659 |
| Astro-TE NN | 1,410 | 954 | 377 | 111 | 123 | 292 | 74 | 70 | 3,411 |
| Endo NN | 962 | 617 | 267 | 277 | 279 | 226 | 165 | 28 | 2,821 |
| Astro-NT NN | 50 | 321 | 235 | 751 | 317 | 86 | 214 | 11 | 1,985 |
| TH Glut | 17 | 45 | 47 | 11 | 1,561 | 4 | 208 | 1 | 1,894 |
| other IT-ET Glut | 1,195 | 209 | 3 | 11 | 3 | 3 | 2 | 19 | 1,445 |
| Microglia NN | 521 | 305 | 204 | 115 | 112 | 59 | 40 | 15 | 1,371 |
| L2/3 IT CTX Glut | 1,155 | 89 | 1 | 3 | 2 | 8 | 1 | 10 | 1,269 |
| MB Glut | 17 | 50 | 25 | 643 | 35 | 2 | 355 | 2 | 1,129 |
| VLMC NN | 31 | 110 | 16 | 30 | 12 | 918 | 4 | 0 | 1,121 |
| L4/5 IT CTX Glut | 1,031 | 18 | 11 | 1 | 3 | 7 | 2 | 2 | 1,075 |
| OPC NN | 328 | 236 | 165 | 127 | 83 | 50 | 42 | 20 | 1,051 |
| CA3 Glut | 738 | 228 | 11 | 29 | 5 | 9 | 5 | 18 | 1,043 |
| MB GABA | 31 | 86 | 105 | 585 | 9 | 2 | 204 | 2 | 1,024 |
| Pvalb Gaba | 522 | 111 | 56 | 18 | 77 | 2 | 200 | 1 | 987 |
| L6 CT CTX Glut | 821 | 64 | 24 | 1 | 0 | 0 | 1 | 2 | 913 |
| L6 IT CTX Glut | 815 | 36 | 5 | 1 | 0 | 6 | 1 | 5 | 869 |
| L5 IT CTX Glut | 803 | 35 | 7 | 1 | 2 | 1 | 1 | 4 | 854 |
| CNU-HYa Glut | 104 | 479 | 3 | 210 | 3 | 4 | 5 | 6 | 814 |
| DG Glut | 31 | 46 | 6 | 5 | 2 | 3 | 1 | 709 | 803 |

Table S8: **Mouse brain: contrast of marker genes** between the on-target Space Ranger cluster and the off-target clusters.

| Genes | Marker set | Cluster | Cells in | Cells out | Raw | ResolVI | Baseline | LUCID ref. | LUCID ref.-free |
|---|---|---|---|---|---|---|---|---|---|
| 2,000 HVGs | myelin | 2 | 6,053 | 18,651 | 14.6 | 14.5 | 295.7 | 42.2 | 28.9 |
| 2,000 HVGs | meningeal | 9 | 1,234 | 35,260 | 44.1 | 49.8 | 102.3 | 78.6 | 83.4 |
| 2,000 HVGs | thalamic | 6 | 1,464 | 36,458 | 91.5 | 83.1 | 31.1 | 106.6 | 132.6 |
| 2,000 HVGs | dentate | 13 | 878 | 35,630 | 37.2 | 37.8 | 32.5 | 32.5 | 33.1 |
| All genes | myelin | 2 | 6,053 | 30,037 | 23.7 | – | 428.7 | 27.7 | 29.7 |
| All genes | meningeal | 9 | 1,234 | 35,260 | 118.2 | – | 77.0 | 101.5 | 145.0 |
| All genes | thalamic | 6 | 1,464 | 35,088 | 23.7 | – | 14.7 | 27.4 | 29.9 |
| All genes | dentate | 13 | 878 | 4,267 | 15.4 | – | 25.3 | 17.7 | 16.8 |
| All genes | cortical excitatory | 16 | 732 | 15,024 | 13.6 | – | 15.4 | 17.3 | 15.8 |

### Supplementary Note 3 — Human colorectal cancer

Table S9 compares the lineage of the assigned type with the published unsupervised annotation of the same section, which does not use the single-cell reference. The published deconvolution labels use the same reference as LUCID and the baseline, with RCTD on 8 $\mu$m bins, and are therefore less independent; they agree with the published unsupervised annotation for 84.7% of the cells. Adaptation of the reference did not change the agreement. The baseline, whose typing step has no ambient term, is closer to the published labels for tumour, epithelium, B and plasma cells, fibroblasts and smooth muscle; LUCID is closer for T cells, myeloid, endothelial and neuronal cells, and the mean over lineages is 70.6% against 69.1%. Reference-free LUCID found 30 clusters, 20 of them tumour clusters; 81.9% of the cells belong to the majority published lineage of their cluster. Marker contrasts between published lineages are in Table S11. On the HVGs, LUCID raised the contrast of T-cell, fibroblast and B-cell markers and ResolVI left it near raw counts. For epithelial markers ResolVI and the baseline raised the contrast, while LUCID with reference lowered it, because it lowered the share of these markers in the normal epithelium itself (from 1.38% to 1.20% of the counts).

Table S9: **Colorectal cancer: agreement with the published annotation** (% of cells with the published lineage), with the reference adapted to the platform and as given.

| Published lineage | Cells | LUCID, adapted | LUCID, as given | Baseline, adapted | Baseline, as given |
|---|---|---|---|---|---|
| Tumor | 102,662 | 95.2 | 95.2 | 97.9 | 98.0 |
| B cells | 24,038 | 57.6 | 57.0 | 72.5 | 73.0 |
| Fibroblast | 22,460 | 75.5 | 75.5 | 79.2 | 79.7 |
| Myeloid | 18,759 | 58.6 | 58.4 | 51.8 | 53.5 |
| Intestinal Epithelial | 15,064 | 80.8 | 80.7 | 89.8 | 90.0 |
| Endothelial | 9,768 | 56.8 | 56.0 | 52.9 | 51.8 |
| Smooth Muscle | 9,508 | 73.5 | 74.9 | 79.1 | 79.9 |
| T cells | 7,257 | 62.4 | 60.4 | 51.8 | 48.6 |
| Neuronal | 213 | 75.6 | 76.1 | 46.9 | 49.3 |
| All cells | 209,729 | 80.5 | 80.4 | 83.7 | 83.9 |
| Mean over lineages | | 70.6 | 70.5 | 69.1 | 69.3 |
| Same of 33 types as published deconvolution | 208,344 | 65.6 | 67.0 | 74.3 | 74.3 |

Table S10: **Colorectal cancer: mixed flag against the published deconvolution class** of the bin under the cell (singlet, or certain and uncertain doublet).

| Method | Published singlets | Flagged (%) | Published doublets | Flagged (%) |
|---|---|---|---|---|
| LUCID, reference | 143,257 | 31.1 | 53,552 | 58.7 |
| LUCID, reference-free | 143,257 | 27.5 | 53,552 | 54.2 |
| Sequential baseline | 143,257 | 46.7 | 53,552 | 91.9 |

### Supplementary Note 4 — Mouse liver

LUCID types per replicate are in Table S12. The published lobule zones give an independent check of the hepatocyte signal: the share of pericentral markers (*Glul, Cyp2e1, Oat, Cyp1a2, Slc1a2, Lect2, Cyp7a1*) and periportal markers (*Cyp2f2, Hal, Sds, Pck1, Ass1, Arg1, Hsd17b13, Gls2*) by zone is unchanged by LUCID (Table S13; ratio between zone 1 and zone 6 for pericentral markers 41 in raw counts, 42 with reference, 42 reference-free). Lineage markers were evaluated in cells with at least three raw counts of the markers of one lineage (Table S14). ResolVI lowered the lineage markers in the cells that carry them and added them to cells that have none.

Without a reference, five of the seven LUCID clusters of the liver followed the lobule zones and two (3,529 cells) gathered the non-parenchymal types.

### Supplementary Note 5 — Platform adaptation and calibration

The gene-wise platform factor was narrow for the tumour, whose reference comes from the same study, and wide for the two atlas references (Fig. S4A; standard deviation of the log factor 0.39, 0.88 and 0.92). Adaptation mattered where the mismatch was large and the tissue is heterogeneous: in the brain the counts kept rose from 31.4% to 55.4% and the median ambient fraction fell from 0.73 to 0.50 (Table S15). In the liver the identifiability rule already protected the hepatocytes and adaptation changed little.

#### *Calibration on control populations*

On real data no population is known to be free of mixed profiles. We took as controls the populations where mixing with another type is least likely (Table S16). The share of cells above the nominal threshold was close to 5% in the liver and well above it in the tumour core, where part of the excess may be real, because tumour cells are in contact with stromal and immune cells.

Table S11: **Colorectal cancer: contrast of lineage markers** between the published lineage where they are expected and the lineages where raw counts show less than a tenth of that share. ResolVI returns only the HVGs.

| Genes | Marker set | Published lineage | Cells in | Cells out | Raw | ResolVI | Baseline | LUCID ref. | LUCID ref.-free |
|---|---|---|---|---|---|---|---|---|---|
| 2,000 HVGs | epithelial | Intestinal Epithelial | 15,064 | 97,336 | 46.5 | 94.3 | 55.7 | 37.8 | 52.8 |
| 2,000 HVGs | smooth muscle | Smooth Muscle | 9,508 | 205,554 | 25.5 | 24.3 | 37.1 | 31.2 | 34.1 |
| 2,000 HVGs | plasma and B | B cells | 24,038 | 168,703 | 60.1 | 63.7 | 45.2 | 75.5 | 79.7 |
| 2,000 HVGs | fibroblast | Fibroblast | 22,460 | 117,726 | 10.9 | 10.4 | 20.2 | 19.7 | 22.9 |
| 2,000 HVGs | T cell | T cells | 7,257 | 127,234 | 10.9 | 10.3 | 15.9 | 19.6 | 21.0 |
| 2,000 HVGs | myeloid | Myeloid | 18,759 | 0 | nan | nan | nan | nan | nan |
| All genes | epithelial | Intestinal Epithelial | 15,064 | 73,031 | 15.6 | – | 18.4 | 28.2 | 30.9 |
| All genes | smooth muscle | Smooth Muscle | 9,508 | 183,094 | 51.8 | – | 100.7 | 56.9 | 58.7 |
| All genes | plasma and B | B cells | 24,038 | 178,221 | 34.3 | – | 24.4 | 47.2 | 48.3 |
| All genes | fibroblast | Fibroblast | 22,460 | 117,726 | 60.8 | – | 146.3 | 63.3 | 72.7 |
| All genes | T cell | T cells | 7,257 | 117,726 | 39.6 | – | 61.0 | 36.5 | 39.4 |
| All genes | myeloid | Myeloid | 18,759 | 117,726 | 23.6 | – | 41.2 | 24.7 | 25.9 |

Table S12: **Mouse liver: cells per LUCID type (with reference).**

| Type | Replicate 1 | Replicate 2 | Total | Share (%) |
|---|---|---|---|---|
| hepatocyte | 18,947 | 26,344 | 45,291 | 91.81 |
| endothelial cell of hepatic sinusoid | 944 | 1,073 | 2,017 | 4.09 |
| hepatic stellate cell | 688 | 906 | 1,594 | 3.23 |
| Kupffer cell | 160 | 83 | 243 | 0.49 |
| natural killer cell | 55 | 23 | 78 | 0.16 |
| myeloid leukocyte | 29 | 15 | 44 | 0.09 |
| B cell | 32 | 9 | 41 | 0.08 |
| plasmacytoid dendritic cell | 18 | 5 | 23 | 0.05 |

Table S13: **Mouse liver: mean share of zonation-marker counts per cell (%)** by published lobule zone, all genes. Cells per zone: 2,732, 10,045, 7,101, 10,642, 16,339, 2,469.

| Markers | Method | Zone 1 | 2 | 3 | 4 | 5 | Zone 6 |
|---|---|---|---|---|---|---|---|
| pericentral | Raw counts | 1.60 | 0.96 | 0.53 | 0.21 | 0.06 | 0.04 |
| pericentral | Sequential baseline | 2.35 | 1.26 | 0.62 | 0.21 | 0.05 | 0.03 |
| pericentral | LUCID, reference | 1.55 | 0.93 | 0.52 | 0.21 | 0.06 | 0.04 |
| pericentral | LUCID, reference-free | 1.62 | 0.96 | 0.52 | 0.21 | 0.06 | 0.04 |
| periportal | Raw counts | 0.37 | 0.65 | 0.84 | 1.18 | 1.62 | 1.41 |
| periportal | Sequential baseline | 0.41 | 0.66 | 0.88 | 1.18 | 1.49 | 1.56 |
| periportal | LUCID, reference | 0.35 | 0.63 | 0.81 | 1.15 | 1.57 | 1.36 |
| periportal | LUCID, reference-free | 0.36 | 0.64 | 0.84 | 1.17 | 1.62 | 1.41 |

Table S14: **Mouse liver: mean share of lineage-marker counts per cell (%)** in cells with at least three raw counts of the lineage markers and in cells with none.

| Genes | Markers and cells | Cells | Raw | ResolVI | Baseline | LUCID ref. | LUCID ref.-free |
|---|---|---|---|---|---|---|---|
| 2,000 HVGs | immune | 4,100 | 0.77 | 0.42 | 0.01 | 1.74 | 0.77 |
| 2,000 HVGs | endothelial | 543 | 0.71 | 0.13 | 0.01 | 0.74 | 0.73 |
| 2,000 HVGs | stellate | 557 | 0.71 | 0.12 | 0.02 | 1.13 | 0.71 |
| 2,000 HVGs | immune, cells without markers | 18,667 | 0.00 | 0.20 | 0.00 | 0.00 | 0.00 |
| 2,000 HVGs | endothelial, cells without markers | 18,667 | 0.00 | 0.09 | 0.00 | 0.00 | 0.00 |
| 2,000 HVGs | stellate, cells without markers | 18,667 | 0.00 | 0.07 | 0.00 | 0.00 | 0.00 |
| All genes | immune | 4,100 | 0.21 | – | 0.00 | 0.81 | 0.22 |
| All genes | endothelial | 543 | 0.19 | – | 0.00 | 0.27 | 0.20 |
| All genes | stellate | 557 | 0.19 | – | 0.01 | 0.45 | 0.19 |
| All genes | immune, cells without markers | 18,667 | 0.00 | – | 0.00 | 0.00 | 0.00 |
| All genes | endothelial, cells without markers | 18,667 | 0.00 | – | 0.00 | 0.00 | 0.00 |
| All genes | stellate, cells without markers | 18,667 | 0.00 | – | 0.00 | 0.00 | 0.00 |

Table S15: **Effect of the platform adaptation** on LUCID with reference (% of counts kept, % of cells flagged as mixed, median ambient fraction).

| Section | sd of log factor | Kept, as given | Kept, adapted | Mixed, as given | Mixed, adapted | Median $\alpha$, as given | Median $\alpha$, adapted |
|---|---|---|---|---|---|---|---|
| Mouse brain | 0.92 | 31.4 | 55.4 | 52.2 | 43.6 | 0.73 | 0.50 |
| Colorectal cancer | 0.39 | 73.5 | 73.5 | 38.9 | 39.4 | 0.19 | 0.18 |
| Mouse liver, replicate 1 | 0.88 | 96.6 | 94.3 | 15.2 | 14.6 | 0.00 | 0.00 |
| Mouse liver, replicate 2 | 0.88 | 97.2 | 94.6 | 16.7 | 15.4 | 0.00 | 0.00 |

Table S16: **Share of control cells above the bootstrap thresholds.**

| Dataset | Mode | Control population | Cells | Above 5% (%) | Above 1% (%) |
|---|---|---|---|---|---|
| Mouse liver | LUCID, reference | hepatocytes without lineage-marker counts | 17,923 | 6.7 | 0.3 |
| Mouse brain | LUCID, reference-free | main cluster of the white-matter niche | 3,551 | 8.1 | 3.3 |
| Colorectal cancer | LUCID, reference | Tumor III cells in the published tumour region | 85,971 | 22.7 | 13.9 |
| Colorectal cancer | LUCID, reference-free | a tumour cluster in the published tumour region | 13,807 | 15.8 | 8.3 |

### Supplementary Figures

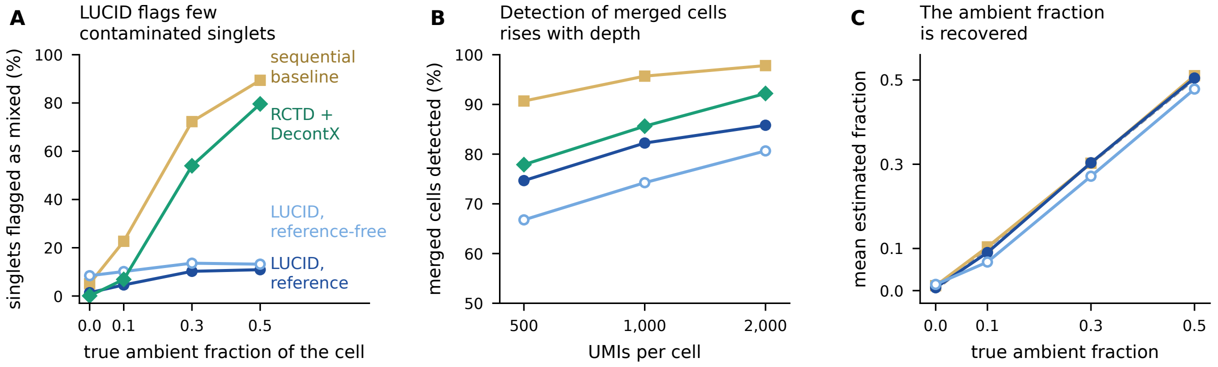


Figure S1: **Simulation with regional contamination** (36,000 cells). (**A**) True singlets flagged as mixed by true ambient fraction. (**B**) True doublets flagged, by counts per cell. (**C**) Mean estimated ambient fraction of the singlets against the true value; dashed line, identity. LUCID at $p < 0.05$.

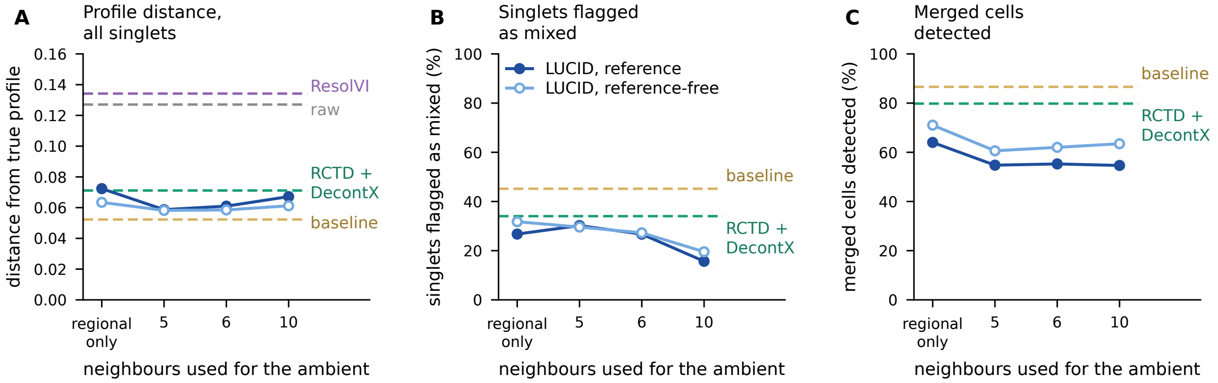


Figure S2: **Spatial simulation: number of neighbours used for the ambient profile.** (**A**) Distance from the true profile on the HVGs, all singlets. (**B**) True singlets flagged as mixed. (**C**) Merged cells flagged. Dashed lines: methods that do not use neighbours.

### Supplementary Note 6 — Comparison with RCTD, DecontX and semi-supervised ResolVI

#### *Pipeline*

To compare LUCID with the published tools that a two-step analysis would use, we ran RCTD (Cable *et al.*, 2022) (spacexr 1.2.0, Bioconductor) in doublet mode on the segmented cells of each section and of the two simulations, followed by DecontX (Yang *et al.*, 2020) (celda 1.26.0) with the first RCTD type of each cell as cluster label. RCTD received the single-cell reference itself, up to 1,000 cells per type from the same atlases used by LUCID, because it estimates its own platform effect. Cells with at least 100 counts were used (`UMI_min = 100`), with otherwise default parameters. Cells called *doublet_certain* or *doublet_uncertain* count as flagged. DecontX was run in two modes: with default settings, where the background is estimated from the cells themselves, and with the counts of the bins not assigned to any cell as background (per niche in the brain and the tumour, per replicate in the liver, with the replicate as batch). Decontaminated counts were rounded to integers. The main text and Fig. 1 use the default mode.

RCTD took 7.3, 26.8 and 0.5 h on the brain, tumour and liver sections with 32, 48 and 8 worker processes (Table S17). These runs used the default multithreaded linear algebra inside each worker, which oversubscribed the nodes, so they are an upper bound. With one thread per worker, each quarter of the tumour section took between 1.7 and 3.2 h on 12 workers.

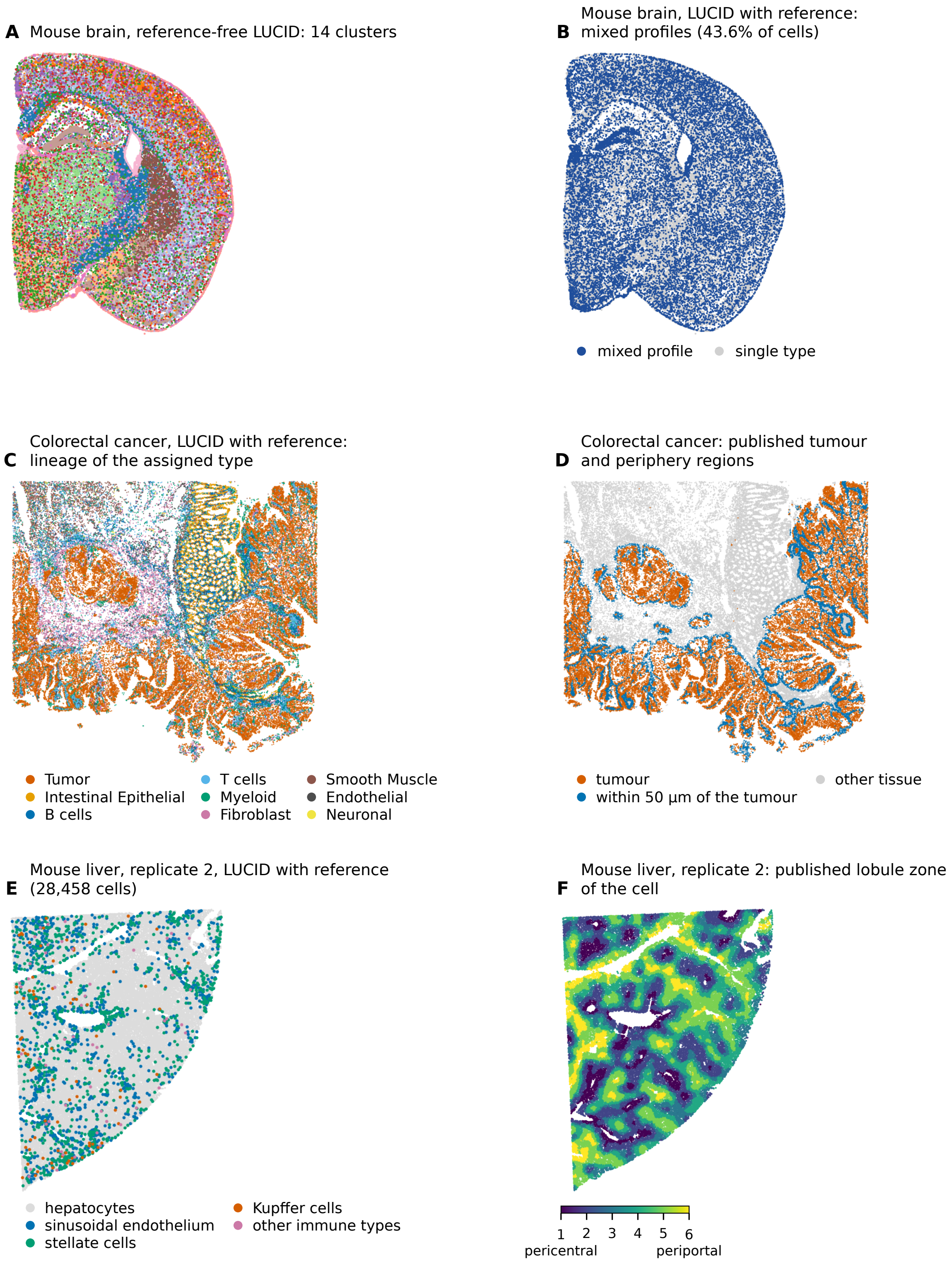


Figure S3: **Maps of the public sections.** (**A**) Mouse brain, reference-free clusters. (**B**) Mouse brain, cells flagged as mixed with reference. (**C**) Colorectal cancer, lineage of the LUCID type (80,000 cells drawn at random). (**D**) Published tumour and periphery regions of the same cells. (**E**) Mouse liver, replicate 2, LUCID types. (**F**) Published lobule zone.

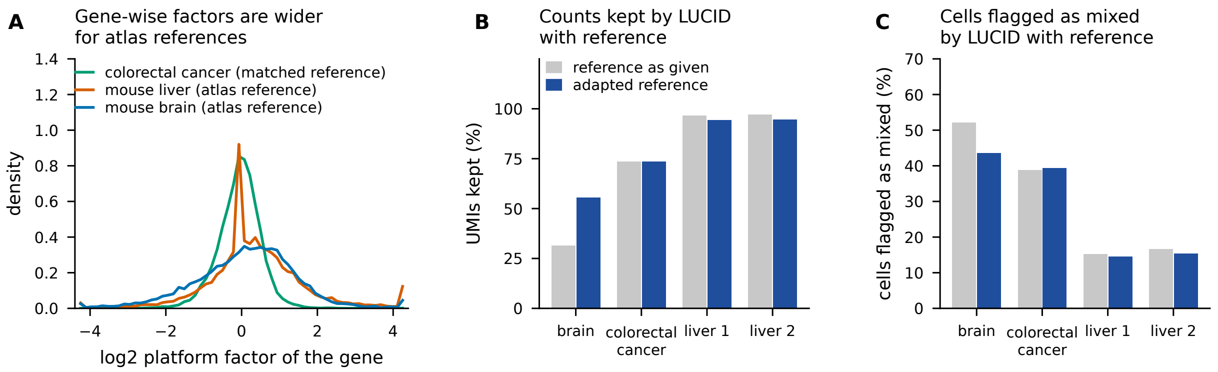


Figure S4: **Platform adaptation of the reference.** (**A**) Gene-wise platform factor of the genes with a pooled share above $10^{-6}$; in the liver, the peak at zero is made of genes absent from the reference. (**B**) Counts kept and (**C**) cells flagged as mixed by LUCID with the reference as given and adapted.

Table S17: **RCTD and DecontX on the three sections.** RCTD classes, share of cells flagged as doublets (certain or uncertain), share of counts kept and cells emptied by DecontX with default settings (first pair) and with the unassigned-bin background (second pair), and RCTD run time.

| Section | Cells | Singlet | Certain | Uncertain | Reject | Doublets (%) | Kept (%) | Emptied | Kept (%) | Emptied | RCTD, h (workers) |
|---|---|---|---|---|---|---|---|---|---|---|---|
| Mouse brain | 39,854 | 6,909 | 18,308 | 12,345 | 2,292 | 76.9 | 81.2 | 458 | 81.9 | 753 | 7.3 (32) |
| Colorectal cancer | 215,405 | 139,308 | 44,615 | 23,896 | 7,586 | 31.8 | 95.3 | 23 | 92.7 | 548 | 26.8 (48) |
| Mouse liver | 49,331 | 39,410 | 6,724 | 3,192 | 5 | 20.1 | 99.3 | 0 | 29.6 | 11,646 | 0.5 (8) |

#### *Simulations*

In the simulation with contamination from the neighbours, RCTD assigned the correct type to 91.8% of the singlets (LUCID with reference 97.0%). It flagged 34.0% of the singlets, almost none of the uncontaminated ones (0.0%) but 54.3% and 74.4% of those with an ambient fraction of 0.3 and 0.5 (Table S18). RCTD therefore reads contamination as a second cell, as the sequential baseline does, while LUCID flags fewer heavily contaminated singlets. In the regional simulation, where a merged cell and a contaminated cell differ, RCTD flagged 35.3% of the singlets against 6.8% for LUCID, with a precision of 0.30 against 0.68. Its detection of merged cells rose with depth (77.9%, 85.6% and 92.2% at 500, 1,000 and 2,000 counts).

After DecontX the distance from the true profile on the HVGs was 0.071 over all singlets and 0.124 for the contaminated cells in mixed neighbourhoods, between raw counts and the methods with an explicit neighbour or regional ambient term (Table S19). With the unassigned-bin background the distances were 0.064 and 0.110. DecontX changed the uncontaminated cells little (0.070 against 0.068 for raw counts).

#### *Mouse brain*

RCTD flagged 76.9% of the brain cells as doublets (Table S17), consistent with a reference from another technology and with contamination read as a second cell. DecontX kept 81.2% of the counts and emptied 458 cells. On the HVGs it raised the contrast of the four marker sets relative to raw counts (myelin 1.92, meninges 2.00, thalamus 1.54, dentate gyrus 1.27). On all genes it lowered the myelin (0.44) and meningeal (0.80) contrasts.

#### *Human colorectal cancer*

The lineage of the first RCTD type agreed with the published unsupervised annotation for 82.0% of the cells, between LUCID (80.5%) and the baseline (83.7%), with a macro recall of 69.1% (LUCID 70.6%) (Tables S21 and S22). RCTD agreed best with the published deconvolution (81.3% of exact types), but that annotation was itself obtained with RCTD in doublet mode on 8 $\mu$m bins (Oliveira *et al.*, 2025), so the comparison is partly circular. RCTD recovered epithelial, stromal and endothelial lineages more often

Table S18: **Typing and flags in the two simulations** (%; precision as a fraction). Singlets flagged by true ambient fraction in the four central columns. LUCID at $p < 0.05$ and six neighbours.

| Method | Type acc. | Singlets fl. | $\alpha = 0$ | 0.1 | 0.3 | 0.5 | Merged det. | Precision |
|---|---|---|---|---|---|---|---|---|
| *Contamination from the neighbours* | | | | | | | | |
| Sequential baseline | 95.3 | 45.2 | 6.1 | 22.9 | 67.5 | 84.5 | 86.6 | 0.25 |
| LUCID, reference | 97.0 | 26.7 | 2.9 | 13.1 | 39.6 | 51.4 | 55.3 | 0.27 |
| LUCID, reference-free | 72.5 | 27.3 | 9.0 | 12.2 | 35.8 | 52.2 | 62.0 | 0.29 |
| RCTD | 91.8 | 34.0 | 0.0 | 7.7 | 54.3 | 74.4 | 79.8 | 0.29 |
| *Regional contamination* | | | | | | | | |
| Sequential baseline | 95.4 | 47.6 | 5.1 | 22.7 | 72.3 | 89.4 | 94.7 | 0.26 |
| LUCID, reference | 99.9 | 6.8 | 1.4 | 4.6 | 10.2 | 10.9 | 80.9 | 0.68 |
| LUCID, reference-free | 73.7 | 11.4 | 8.4 | 10.1 | 13.6 | 13.2 | 73.8 | 0.54 |
| RCTD | 94.7 | 35.3 | 0.1 | 6.9 | 54.0 | 79.6 | 85.2 | 0.30 |

Table S19: **Distance from the true profile in the simulation with contamination from the neighbours**, on the 2,000 HVGs (first three columns) and on all genes (last three). Mixed: ambient fraction of at least 0.3 and at least half of the neighbours of other types. ResolVI returns only the HVGs.

| Method | All | Uncont. | Mixed | All | Uncont. | Mixed |
|---|---|---|---|---|---|---|
| Raw counts | 0.127 | 0.068 | 0.233 | 0.080 | 0.086 | 0.140 |
| ResolVI | 0.134 | 0.075 | 0.214 | – | – | – |
| Sequential baseline | 0.052 | 0.068 | 0.093 | 0.064 | 0.086 | 0.116 |
| RCTD + DecontX | 0.071 | 0.070 | 0.124 | 0.078 | 0.087 | 0.138 |
| RCTD + DecontX, background | 0.064 | 0.070 | 0.110 | 0.073 | 0.087 | 0.131 |
| LUCID, reference | 0.061 | 0.070 | 0.094 | 0.081 | 0.087 | 0.133 |
| LUCID, reference-free | 0.058 | 0.072 | 0.093 | 0.069 | 0.088 | 0.114 |

than LUCID and T cells much less often (24.6% against 62.4%). On the HVGs, RCTD + DecontX raised the contrast of the immune and stromal markers between their published lineage and the others more than LUCID (Table S20): 22.0 against 19.6 for T-cell markers and 111.5 against 19.7 for fibroblast markers. Both lowered the contrast of epithelial markers (35.2 and 37.8, raw counts 46.5).

Table S20: **Colorectal cancer: contrast of marker genes** between the published unsupervised lineage where they are expected and the lineages where raw counts show less than a tenth of that share, on the 2,000 HVGs.

| Method | T cell | Plasma and B | Fibroblast | Smooth muscle | Epithelial |
|---|---|---|---|---|---|
| Raw counts | 10.9 | 60.1 | 10.9 | 25.5 | 46.5 |
| ResolVI | 10.3 | 63.7 | 10.4 | 24.3 | 94.3 |
| ResolVI, semi-supervised | 11.2 | 63.5 | 11.5 | 25.8 | 78.1 |
| Sequential baseline | 15.9 | 45.2 | 20.2 | 37.1 | 55.7 |
| RCTD + DecontX | 22.0 | 98.8 | 111.5 | 36.0 | 35.2 |
| RCTD + DecontX, background | 12.0 | 67.9 | 147.0 | 49.5 | 70.1 |
| LUCID, reference | 19.6 | 75.5 | 19.7 | 31.2 | 37.8 |
| LUCID, reference-free | 21.0 | 79.7 | 22.9 | 34.1 | 52.8 |

***Mouse liver***

RCTD assigned 49,270 of the 49,331 liver cells (99.9%) to hepatocytes, 30 to stellate cells and 18 to sinusoidal endothelium, so DecontX received a single cluster in practice. With default settings it estimated a median contamination of 0.022, kept 99.3% of the counts and lowered the immune and stellate marker shares in the cells that carry them (Table S23). With the unassigned-bin background, whose profile is

Table S21: **Colorectal cancer: agreement with the published annotations** (%). Lineage of the first RCTD type or of the LUCID and baseline type compared with the published unsupervised annotation (unsup., also as macro recall over lineages) and with the published deconvolution (deconv., exact type and lineage).

| Method | Lineage, unsup. | Macro recall | Type, deconv. | Lineage, deconv. |
|---|---|---|---|---|
| Sequential baseline | 83.7 | 69.1 | 74.3 | 82.1 |
| RCTD | 82.0 | 69.1 | 81.3 | 85.7 |
| LUCID, reference | 80.5 | 70.6 | 65.6 | 79.5 |

Table S22: **Colorectal cancer: recall by published unsupervised lineage** (%).

| Published lineage | Cells | Baseline | RCTD | LUCID, ref. |
|---|---|---|---|---|
| Tumor | 102,662 | 97.9 | 97.9 | 95.2 |
| B cells | 24,038 | 72.5 | 47.2 | 57.6 |
| Fibroblast | 22,460 | 79.2 | 84.0 | 75.5 |
| Myeloid | 18,759 | 51.8 | 55.3 | 58.6 |
| Intestinal Epithelial | 15,064 | 89.8 | 95.4 | 80.8 |
| Endothelial | 9,768 | 52.9 | 66.1 | 56.8 |
| Smooth Muscle | 9,508 | 79.1 | 84.9 | 73.5 |
| T cells | 7,257 | 51.8 | 24.6 | 62.4 |
| Neuronal | 213 | 46.9 | 66.2 | 75.6 |

close to the hepatocyte profile, the median contamination rose to 0.95: DecontX kept 29.6% of the counts, emptied 11,646 cells and inflated the pericentral zonation ratio from 40.5 to 807.0. This is the identifiability problem that the rule of LUCID addresses: when the ambient profile equals the cell's own profile, the ambient fraction cannot be estimated from the data.

Table S23: **Mouse liver: lineage markers and zonation.** Mean share of lineage-marker counts (%) in cells with at least three immune, endothelial or stellate marker counts in the raw data, on the HVGs (first three columns) and on all genes (next three). Zonation: ratio of the mean share of pericentral markers between zone 1 and zone 6, and of periportal markers between zone 6 and zone 1, on all genes.

| Method | Immune | Endoth. | Stellate | Immune | Endoth. | Stellate | Pericentral | Periportal |
|---|---|---|---|---|---|---|---|---|
| Raw counts | 0.77 | 0.71 | 0.71 | 0.21 | 0.19 | 0.19 | 40.5 | 3.8 |
| ResolVI | 0.42 | 0.13 | 0.12 | – | – | – | – | – |
| Sequential baseline | 0.01 | 0.01 | 0.02 | 0.00 | 0.00 | 0.01 | 88.2 | 3.8 |
| RCTD + DecontX | 0.64 | 0.70 | 0.57 | 0.17 | 0.18 | 0.15 | 40.3 | 3.8 |
| RCTD + DecontX, background | 0.14 | 0.44 | 0.54 | 0.03 | 0.11 | 0.14 | 807.0 | 0.3 |
| LUCID, reference | 1.74 | 0.74 | 1.13 | 0.81 | 0.27 | 0.45 | 41.9 | 3.9 |
| LUCID, reference-free | 0.77 | 0.73 | 0.71 | 0.22 | 0.20 | 0.19 | 42.0 | 3.9 |

***Semi-supervised ResolVI***

ResolVI can use cell-type labels of the spatial cells. With the first RCTD type as label, the distance from the true profile in the simulation fell from 0.134 to 0.114 over all singlets and from 0.214 to 0.178 for the contaminated cells in mixed neighbourhoods, still above RCTD + DecontX and LUCID (Table S24). On the real sections the labels changed little: the brain and tumour contrasts stayed close to raw counts, and in the liver, where RCTD labelled almost all cells as hepatocytes, the lineage-marker shares and the flattened zonation were those of the unsupervised run. Semi-supervised ResolVI is shown next to the unsupervised run in Fig. 1B, F and H.

Table S24: **ResolVI without and with labels**, compared with raw counts, RCTD + DecontX and LUCID with reference. Simulation with contamination from the neighbours, distances on the HVGs. Contrasts and marker shares on the HVGs as in Fig. 1F–H. Pericentral zonation: ratio of the mean share of pericentral markers between zone 1 and zone 6, on the HVGs.

| | Raw | ResolVI | ResolVI, labels | RCTD + DecontX | LUCID, ref. |
|---|---|---|---|---|---|
| *Simulation: distance from the true profile* | | | | | |
| All singlets | 0.127 | 0.134 | 0.114 | 0.071 | 0.061 |
| Uncontaminated | 0.068 | 0.075 | 0.069 | 0.070 | 0.070 |
| Contaminated, mixed | 0.233 | 0.214 | 0.178 | 0.124 | 0.094 |
| *Mouse brain: contrast relative to raw counts* | | | | | |
| Myelin | 1.00 | 0.99 | 1.17 | 1.92 | 2.89 |
| Meninges | 1.00 | 1.13 | 1.07 | 2.00 | 1.78 |
| Thalamus | 1.00 | 0.91 | 0.90 | 1.54 | 1.16 |
| Dentate gyrus | 1.00 | 1.01 | 1.04 | 1.27 | 0.87 |
| *Colorectal cancer: contrast* | | | | | |
| T cell | 10.9 | 10.3 | 11.2 | 22.0 | 19.6 |
| Plasma and B | 60.1 | 63.7 | 63.5 | 98.8 | 75.5 |
| Fibroblast | 10.9 | 10.4 | 11.5 | 111.5 | 19.7 |
| *Mouse liver: share of lineage-marker counts (%)* | | | | | |
| Immune | 0.77 | 0.42 | 0.38 | 0.64 | 1.74 |
| Endothelial | 0.71 | 0.13 | 0.12 | 0.70 | 0.74 |
| Stellate | 0.71 | 0.12 | 0.12 | 0.57 | 1.13 |
| Pericentral zonation | 56.5 | 34.6 | 35.4 | 56.4 | 58.3 |
